# Cortical cellular encoding of thermotactile integration

**DOI:** 10.1101/2023.04.23.537915

**Authors:** Philipp Schnepel, Ricardo Paricio-Montesinos, Ivan Ezquerra-Romano, Patrick Haggard, James FA Poulet

## Abstract

Recent evidence suggests that primary sensory cortical regions play a role in the integration of information from multiple sensory modalities. How primary cortical neurons integrate multisensory information is unclear, partly because multisensory interactions in the cortex are typically weak or modulatory. To address this question, we take advantage of the robust representation of thermal (cooling) and tactile stimuli in mouse forepaw primary somatosensory cortex (fS1). Using a thermotactile detection task, we show that the perception of threshold level cool or tactile information is enhanced when they are presented simultaneously compared to presentation alone. To investigate the cortical correlates of thermotactile integration, we performed in vivo extracellular recordings from fS1 during unimodal and bimodal stimulation of the forepaw. Unimodal stimulation evoked thermal- or tactile- specific excitatory and inhibitory responses of fS1 neurons. The most prominent features of bimodal, thermotactile stimulation are the recruitment of unimodally silent fS1 neurons, non-linear integration features and a change in the response dynamics to favor longer response durations. Together, we identify quantitative and qualitative changes in cortical encoding that may underlie the improvement in perception of multisensory, thermotactile surfaces during haptic exploration.

## Introduction

A fundamental function of the brain is the integration of different streams of sensory input. Quantitative behavioral testing in humans and animal models has shown that multisensory integration enhances perceptual performance in different ways ^1–10^. Most of our understanding of the neural encoding of multisensory stimuli, such as the principles of inverse effectiveness or spatial and temporal congruency, comes from studies in the superior colliculus ^11–13^. The neocortex is required for sensory perception, and thought to play a prominent role in multisensory integration ^7^, but far less is known about the rules of multisensory integration in cortical circuits.

Cortical multisensory integration has been thought of as a hierarchical process, with unisensory, primary cortices providing converging input to higher order regions for integration with other modalities ^14–18^. Recently, recordings and anatomical work has suggested that even the earliest stages of cortical sensory processing play a significant role in multisensory integration ^4,19–25^. However, the responses of primary cortical sensory areas to other modalities are typically sparse and weak ^22,26^ and inputs from other modalities have been thought of as modulatory or even mere artefacts of behaviorally generated activity ^27^. Overall, in contrast to the well-defined principles of multisensory integration in the superior colliculus, the cellular representation during multisensory integration in the cortex remains unresolved.

An ideal model to examine cortical mechanisms of multisensory integration is the thermotactile system. Thermal and tactile pathways have dedicated peripheral sensory afferent neurons expressing different ion channel receptors, but everyday perceptual experience tells us that thermal and tactile information are integrated during haptic exploration of object surfaces. Intriguingly, somatosensory illusions also hint at a profound interaction between thermal and tactile pathways even at early stages of the sensory pathway. For example, the lack of dedicated wetness hygroreceptors in primary somatosensory afferent neurons indicates that our sense of wetness is created centrally by the integration of thermal with tactile information ^28^. Further, Weber’s ‘Thaler illusion’ shows that colder objects appear heavier than those of a neutral temperature ^29^, and ‘Thermal referral’ shows that a warm coin placed on the skin next to a cold coin also starts to feel cold ^30–32^.

In previous work, we showed that mouse primary forepaw somatosensory cortex (fS1) contains neurons responsive to cool and touch stimulation of the glabrous skin of the forepaw ^33,34^. However, whether mice show changes in perceptual performance during thermotactile processing, and how fS1 cortical neurons integrate cool and touch inputs has not been examined. The goal of this study was to identify encoding features of cortical neurons during thermotactile integration and examine how they correlate with performance during thermotactile integration. To address this, we developed a high resolution, thermotactile detection task for head-fixed mice that allowed us to deliver separate or combined cool and touch stimuli to the forepaw, and went on to perform single unit recordings from fS1 in awake and anesthetized mice during sensory stimulation.

## Results

### The addition of a second modality enhances cool and touch detection rates and lowers perceptual threshold

Mice use their forepaws with great dexterity during foraging, food consumption, haptic exploration and social interactions. Much like human hands, the mouse forepaw is extremely sensitive to tactile and thermal stimuli and can detect milli-Newton scale forces and temperature changes of < 1 °C ^34–36^. Inspired by a multisensory mouse audio-visual detection task ^37^, we designed a thermotactile Go/NoGo detection task to assess the psychophysical abilities of mice to detect uni- and multi-modal thermotactile stimuli (Fig. 1A). We hypothesized that combining cool with touch stimuli would enhance perceptual performance as compared to presentation of a unimodal stimulus alone.

**Figure 1:**
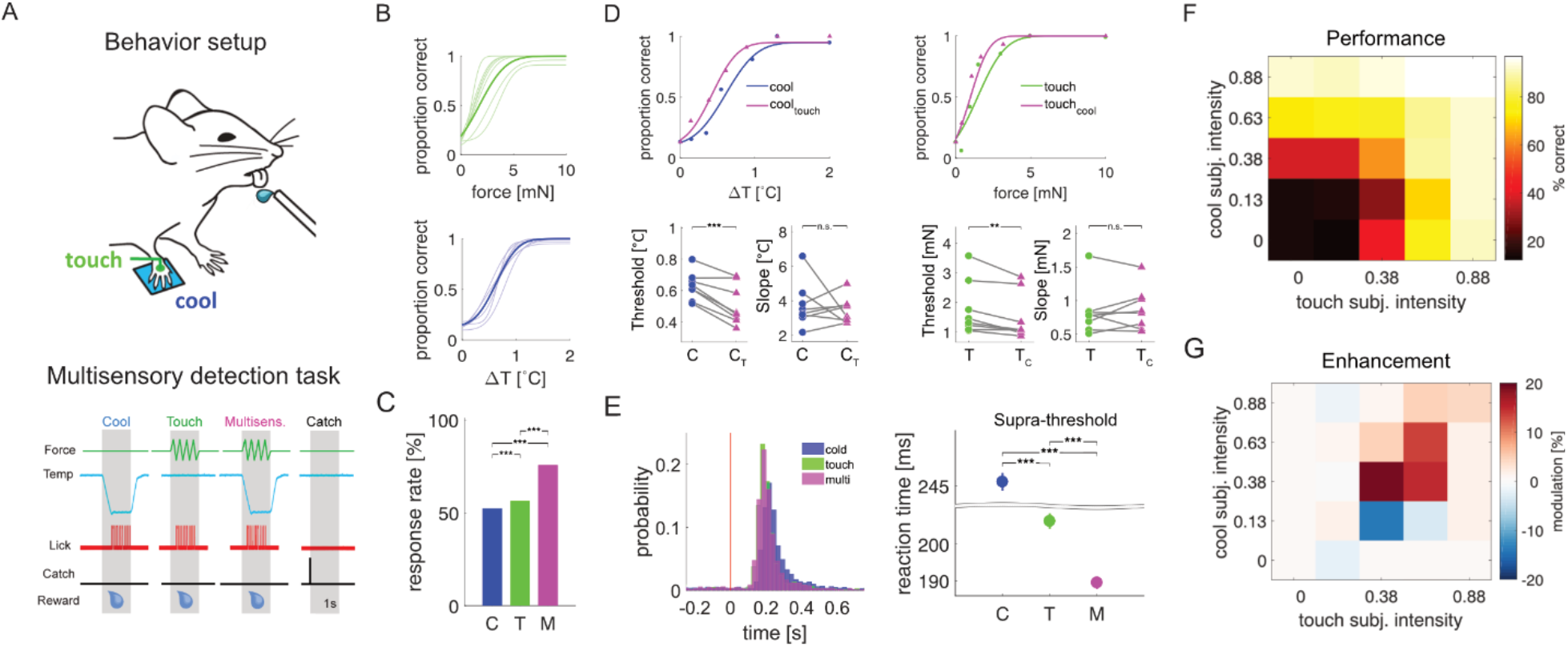
Mice exhibit performance enhancement in a thermo-tactile detection task. A) Behavioral setup and detection task paradigm. Mice are head-fixed and receive tactile and temperature stimuli to their right forepaw. Go/No-go task design with rewarded (stimulus) and unrewarded (catch) trials. B) Fitted psychometric curves for touch and cool stimuli for all mice (right, n=8, average in bold). C) Average response rates across all animals, significant differences between means are denoted by stars (paired t-test). D) Fitted psychometric curves for cool (blue) and cool + subthreshold touch (magenta, left panel) and touch (green) and touch + subthreshold cool (magenta, right panel) for an example animal. Thresholds and slopes from the fitted psychometric functions for each animal are shown on the bottom. Significant differences between means are denoted by stars (paired t-test). E) Average first lick latency (‘reaction time’) for supra-threshold cool, touch and bimodal stimuli across all mice. Error bars are mean ± SEM, significant differences between means are denoted by stars (paired t-test). F) Normalized % correct response rates for all stimulus combinations (‘subjective intensity’), averaged across all animals. G) Relative multisensory enhancement (modulation: (M-Umax)/Umax) for each stimulus combination reveals strongest enhancement around threshold. Axis tick labels denote bin centers.

Head-restrained, paw-tethered mice were initially trained to report the presence of a cool or touch stimulus by licking a waterspout. Once they reached a stable performance level, we generated session-by-session psychometric curves for cool and touch stimuli with an adaptive staircase procedure (Fig. 1B, Fig. S1A) and pseudo-randomized the delivery of bimodal (cool and touch) and unisensory (cool or touch) stimuli (for details see Methods). For each mouse, perceptual thresholds were combined across 5-11 testing sessions to increase statistical validity (on average 1959 trials per mouse, range: 1138-2944). Average unisensory perceptual thresholds were 0.63 ± 0.09 °C for cool and 1.77 ± 0.9 mN for touch stimuli (n = 8 mice).

Hallmarks of multisensory behavioral enhancement in other systems are lower response latencies and higher stimulus detection rates, when compared to responses to unimodal stimuli. Consistent with this, in our task mice showed higher detection rates to bimodal than unimodal stimuli (Fig. 1C, p < 0.001). Moreover, while mice responded to cool stimuli with longer latency than to touch stimuli (246.1 ± 2.4 ms vs. 206.3 ± 2.1 ms, p = 1.0×10^−34^; Figs 1E, S1B), they responded to thermotactile stimuli with shorter latency than to cool or touch stimuli (189.5 ± 1.7 ms, Fig. 1E). We went on to further assess perceptual performance by fitting psychometric functions to the detection rates to different amplitude uni- and bi-modal stimuli (top graphs Fig. 1D) and compared the threshold for each mouse. Supporting our hypothesis of increased detection performance, both the cool and touch detection thresholds were decreased when adding subthreshold stimuli of the second modality (cool: 0.63 ± 0.09 °C vs. cool+_touch_: 0.51 ± 0.13 °C, p = 9.3×10^−4^; touch: 2.20 ± 1.0 mN vs. touch+_cool_: 1.49 ± 0.79 mN, p = 0.009, bottom graphs Fig 1D).

Detection performance can also be evaluated by the steepness of the slope of the psychometric function around threshold, with a steeper slope indicating higher detection reliability which relates to an increased ‘precision’ of task performance. By comparing the slope between the uni- and bi-modal conditions we observed that across the population there was no overall difference (cool: 3.74 ± 1.31 °C vs. cool+_touch_: 3.44 ± 0.77 °C, p = 0.6 and touch: 0.83 ± 0.36 mN vs. touch+_cool_: 0.87 ± 0.32 mN, p = 0.54), indicating that there was no significant improvement in detection reliability to thermotactile stimuli.

In order to quantify the average amount of multisensory enhancement across all mice, we normalized the stimulus space and calculated the proportion of correct responses, (Fig. 1F,), as well as the difference between the largest unimodal and bimodal response rate (Fig. 1G). These plots show that the degree of multisensory enhancement is largest around threshold (0.5) for both modalities. Taken together, our data show that mice have an enhanced performance during bimodal, thermotactile, detection task as compared to the unimodal detection tasks.

### Cool and touch unimodal stimulation both excite and inhibit fS1 neurons

While we have previously shown that fS1 neurons can respond to cool and touch ^33,34^, the degree of overlap between cool and touch responsiveness has not been characterized at the population level. To address this, and attempt to identify neural correlates of the enhanced multisensory perceptual performance observed in Fig. 1, we went on to perform extracellular recordings from fS1 neurons in awake mice (Fig. 2A, B). To control for the effects of arousal and movement, we also performed recordings under identical conditions, but in isoflurane anaesthetized mice (Fig. S2).

**Figure 2:**
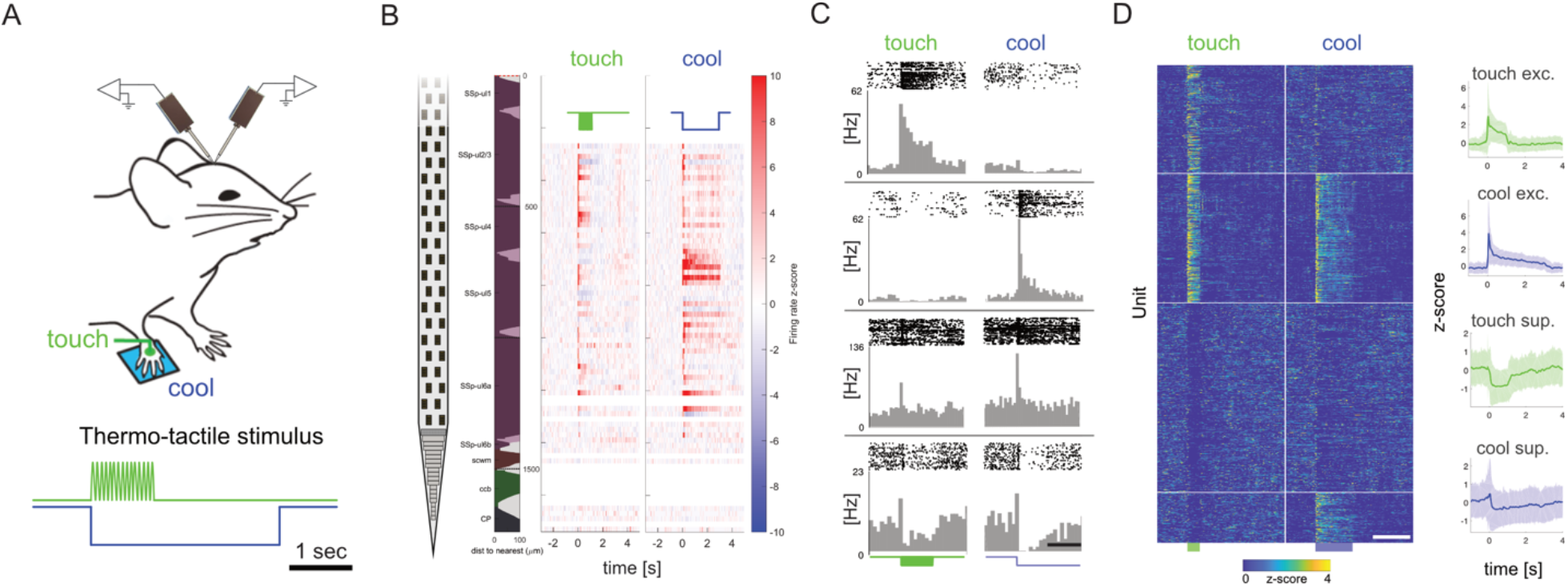
Extracellular recordings of touch and cold representation in forepaw S1. A) Recording setup: Mice are awake, head-fixed and receive tactile (60 Hz vibration, 1 s) and cool (ramp-hold-return, 3 s) stimuli to their right forepaw during extracellular recordings with up to two neuropixel probes. B) Example penetration with a neuropixel probe showing the reconstructed location of probe sites across cortical layers and z-scored firing rate responses across the probe shank for touch and cool stimulation. (SSp-ul: primary somatosensory area upper limb; scwm: supra-callosal cerebral white matter; ccb: corpus callosum body; CP: Caudoputamen). C) Responses of four example units to touch and cool stimulation exhibiting heterogeneous response dynamics to both modalities. Scale bar is 1 s. D) Hierarchical clustering of maximum unimodal stimulus (20 mN / -8 °C) responses of all recorded units (n = 1149). Each row shows the concatenated, z-scored response to maximum touch (left) and cool (right) stimulation for one unit. Units are arranged by cluster to reveal the overlap between touch and cool representations in the population. Scale bar is 3 seconds. On the right, the z-scored and averaged response of touch/cool excited and touch/cool suppressed (or unresponsive) clusters are depicted (shaded area denotes ±2 SD).

Recordings were targeted to left fS1 using intrinsic optical imaging and the recording sites were confirmed post-hoc using DiI staining of the probe tract (Fig. S3). We presented cool, touch or simultaneous cool and touch stimuli to their right forepaw at different amplitudes (5, 10, 20 mN for touch and -1, -2, -4 and -8 °C from a baseline of 32 °C for cool). We first examined single unit responses to unimodal cool and touch stimuli (Fig. 2C) and found different combinations of excitatory and inhibitory responses that we quantified with hierarchical clustering. Clustering to each modality showed that 41% of recorded neurons were unresponsive or inhibited during unisensory stimulation, 32.7% neurons showed excitatory responses to both cooling and touch, 11.3 % were excited by cooling only and 15.1 % to touch only. Of cooling responsive neurons, 33 % were unresponsive or inhibited by touch stimulation. Of the touch responsive neurons, 45 % were unresponsive or inhibited by cool stimulation (Fig. 2D). These results show that fS1 neurons exhibit varying levels of modality-specific excitation and inhibition to thermotactile stimuli.

### Multisensory stimulation boosts cortical responsiveness and recruits previously silent neurons

Prior work in different systems has suggested that responses to multisensory stimuli are amplified as compared to responses to unisensory stimuli ^7^. To examine whether this was the case in the thermotactile system, we concentrated on fS1 neurons that were significantly excited by any stimulus type (cool alone, touch alone, combined cool and touch). Then, we compared their responses to unisensory with multisensory stimuli at a range of stimulus amplitudes (Fig. 3). The median excitatory response strength to the maximum intensity unisensory cool and touch stimuli was comparable (cool 1.63 Hz vs. touch 2.00 Hz, p = 0.98), whereas there was a significant increase in the multimodal response strength (4.75 Hz, p = 6.6e^-16^ and p = 1.2e^-13^ for cool and touch, respectively; Fig. 3A,B,C). Similar to the psychophysical performance, the median fS1 neuron response latency to cool was delayed compared to touch (48 ms cool vs. 21 ms touch, p = 8.8×10^−10^, Fig. 3D), but was longer than the multisensory response latency (28 ms). Similar results were observed in anesthetized mice (Fig. S2A,B), indicating that these results are not due to changes in arousal level or movement. Overall, thermotactile stimulation drives the fS1 circuit more effectively than unisensory cool or touch stimuli.

**Figure 3:**
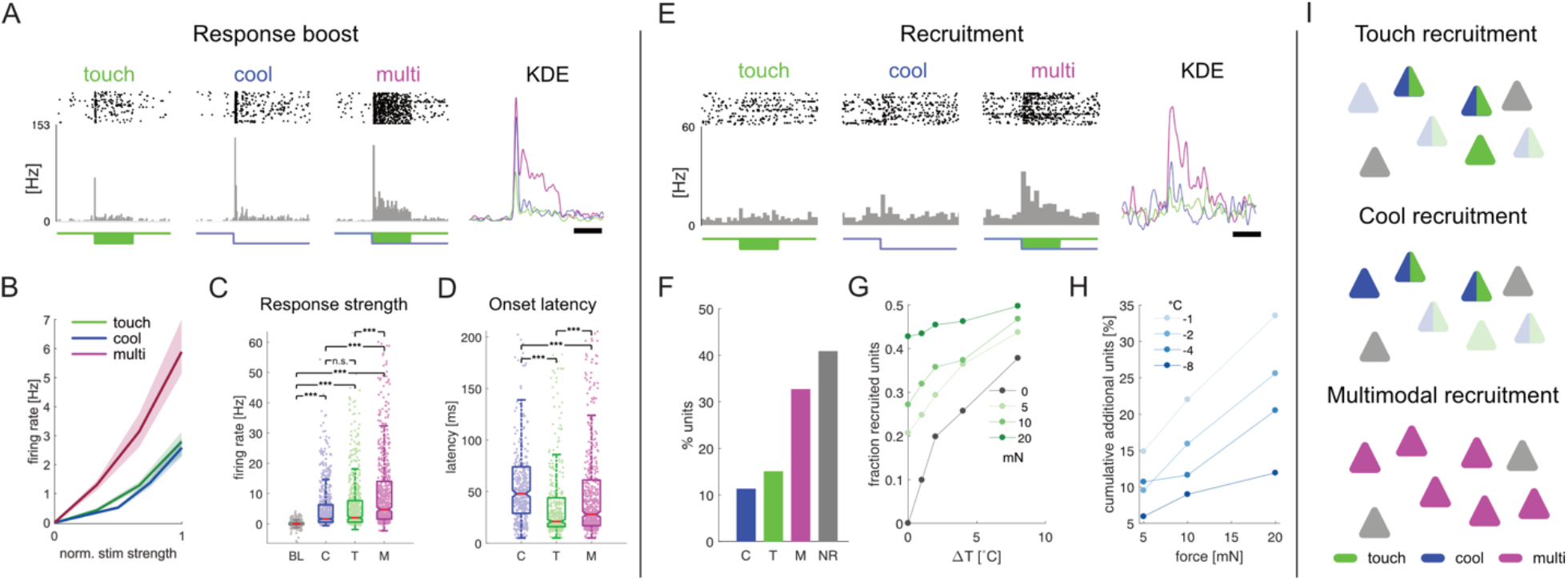
Multimodal stimulation boosts responses and recruits neurons. A) Responses of one example unit to touch, cool and multimodal stimulation. Kernel density estimate (KDE) comparison shows increased response to multimodal stimulation (scale-bar: 500msec). B) Average firing rate of all responsive units to increasing stimulus intensity (normalized) for touch (5,10 and 20 mN), cool (-1,-2,-4 and -8 °C) and multimodal stimulation. C) Maximum, baseline-corrected response strength of all responsive units for each stimulus condition (C=cool, T=touch, M=multimodal; BL=baseline is added for comparison). Significant differences in the medians were determined by multiple comparison testing (Kruskal-Wallis test). D) Same as in C) but for onset latency. E) Responses of one example unit to touch, cool and multimodal stimulation. Kernel density estimate (KDE) comparison shows recruitment in the multimodal condition (scale-bar: 500msec). F) Fractions of responsive (cool, touch, multimodal) and unresponsive single units at maximum intensity stimulation. G) The fraction of recruited units as a function of stimulus intensity illustrates the ceiling effect on the population level. H) The majority of additional recruitment in the multimodal condition happens already at lower intensity stimulation. I) Touch or cool stimulation will recruit a subset of neurons that respond to the respective single modality or to both (gray triangles are unresponsive neurons). Multimodal stimulation will not only recruit all exclusively unimodal neurons but also ‘silent’ neurons that did not reach threshold in the unimodal conditions or were suppressed by modality-specific inhibition (transparent units in the unimodal conditions).

Although we found that many neurons could exhibit excitation for one modality and inhibition for the other (Fig. 2C, D), most responsive neurons were multimodal (Fig. 3F). We reasoned that unimodally silent or inhibited neurons could be unmasked by the combined multimodal drive and thus recruited to increase the numbers of cells representing a multisensory stimulus in fS1 (e.g. unit in Fig. 3E). To quantify this, we went on to examine the dependence of neuron recruitment on stimulus amplitude and calculated the fraction of responsive units at increasing stimulus intensities during unimodal and bimodal stimulation (Fig. 3G). As expected, the number of recruited neurons increases with increasing unimodal cool amplitude (grey line). When including a simultaneous touch stimulus (green lines), the number of recruited neurons increases (as seen by the vertical shift in the green lines at a given thermal stimulus amplitude). This increase in recruitment plateaus towards the maximum touch and cool stimulus strengths, likely due to the network operating at the upper end of its’ dynamic range (‘ceiling effect’). Thus, at low stimulus intensities, adding a second modality results in relatively more units being recruited than at higher stimulus intensities. This can be seen by the steeper increase in the cumulative additional units for weak cooling during touch stimulation than for higher amplitude cooling during touch stimulation (light blue compared to dark blue lines in Fig. 3H). These relationships were also observed in anesthetized recordings, and therefore not due to arousal or movement-related activity (Fig. S2D, E). Together these data suggest that the recruitment of unimodally silent or inhibited neurons is a critical feature of multisensory stimulus representation in fS1 (Fig. 3I)

### Non-linear processing in fS1 during thermotactile integration

A key question in multisensory processing is whether the neural responses to multisensory stimuli are predictable from their responses to unisensory stimuli. A widely used metric to address this is ‘response additivity’. Classically, the response of a unit to multisensory stimulation is compared to the arithmetic sum of the corresponding unimodal responses. Then, the units can be categorized as supra-additive, additive or sub-additive (Fig. 4A, see Methods). To investigate these sub-populations in our dataset, we calculated a ‘Multisensory Index (MSI)’ as a normalized quantification of multimodal enhancement/suppression (see Methods). We observed examples of supra-additive (Fig. 4B, top, MSI > 0), additive (Fig. 4B, middle, MSI ∼ 0) and sub-additive units (Fig. 4B, bottom, MSI < 0) during thermotactile stimulation as compared to unimodal stimulation. The majority of fS1 units were classified as supra- or sub-additive (49.7 and 39.3 %, respectively) and only a smaller fraction as additive (11.0 %, Fig. S5B). Plotting the multimodal responses against the corresponding sum of both unimodal responses (Fig. 4E) showed a clear separation from the isocline for the supra and sub-additive populations, confirming that most thermotactile responses in fS1 are non-linear and not predictable from their unimodal response.

**Figure 4:**
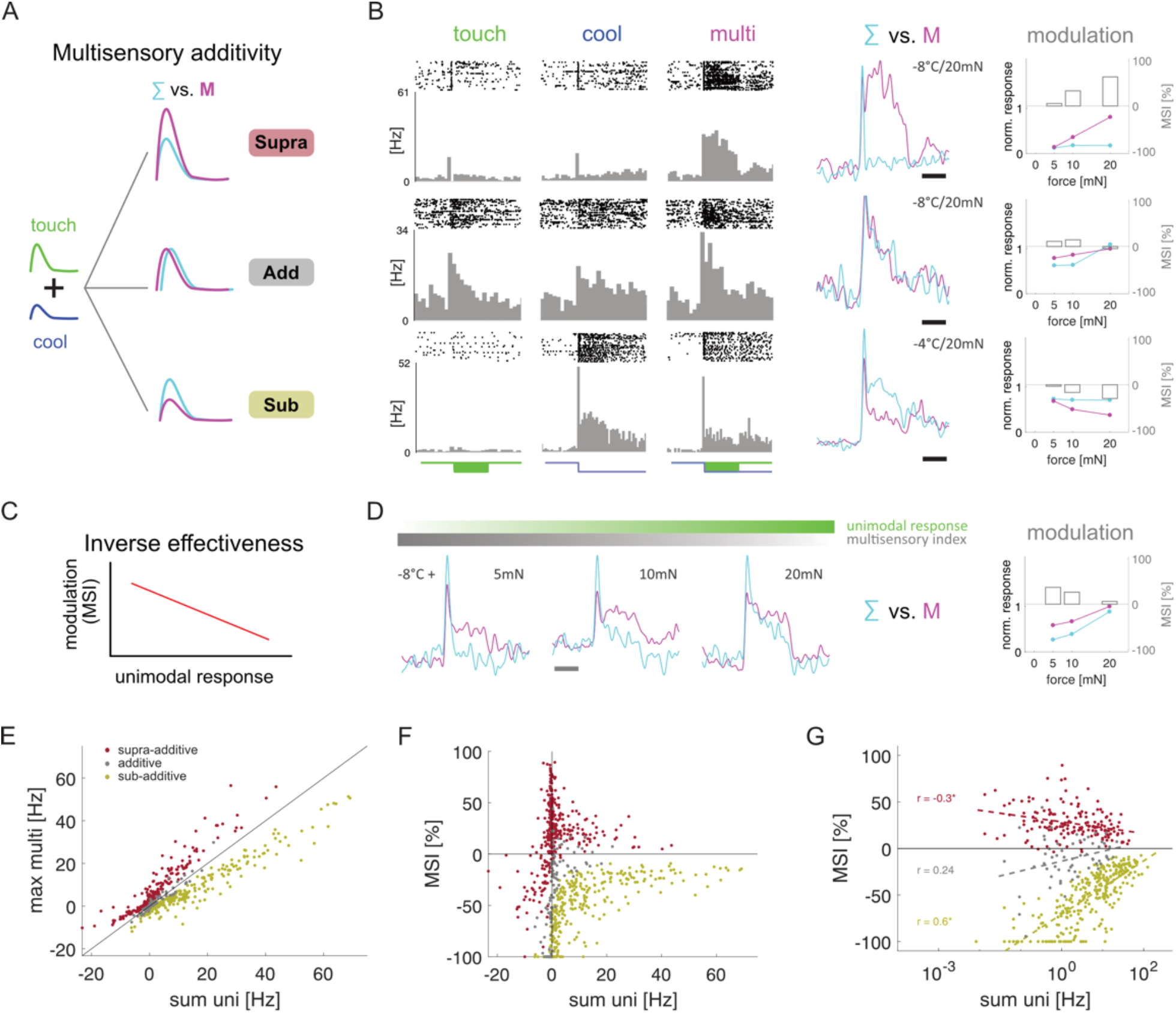
Additivity and inverse effectiveness on the single unit and population level. A) Comparing the arithmetic combination of touch and cool responses (∑) to the response in the multimodal condition (M) allows the definition of additive (gray) and sub-additive (yellow) supra-additive (red) sub-populations of neurons (see Methods). B) Example units illustrating additivity: supra-additive (top), additive (middle) and sub-additive (bottom). Each row shows (from left to right): Raster/PSTH for touch, cool and multimodal stimulus, kernel density estimate (KDE) of the firing rate for ∑ vs. M (scale-bar: 500 ms), normalized response strength for ∑ and M (left axis) and corresponding MSI (gray bars, right axis) for one temperature (the example’s stimulus combination is marked next to the KDE plot, e.g. -4 °C / 20 mN). C) The principle of inverse effectiveness postulates that the strength of the multisensory modulation (MSI, Multisensory Index, see Methods) is inversely correlated to the unimodal stimulus/response strength. D) Example unit illustrating inverse effectiveness. The modulation strength (difference between KDE for ∑ vs. M, gray bar) is decreasing with increased intensity of one modality (tactile strength increase from left to right, green bar; cool is fixed at -8 °C). This is summarized in the modulation plot as in B). E) Maximum multimodal response plotted against the arithmetic sum of the corresponding unimodal stimulus responses for the ‘best’ stimulus combination (see Methods) of each unit (color code as in A). F) MSI plotted against the arithmetic sum of the corresponding unimodal stimulus responses. G) Same as in F but on a semi-log scale. A linear regression of the data reveals a significant negative correlation (i.e. inverse effectiveness) for the supra-additive population, but also a positive correlation for the additive and sub-additive populations.

Classic studies in the superior colliculus have proposed a second potential mechanism to boost low-saliency stimuli and improve signal detection termed ‘inverse effectiveness’. It suggests that a neuron’s multisensory enhancement is inversely proportional to the unisensory response strength (Fig. 4C, example cell in Fig. 4D). As mentioned above, the focus of multimodal integration analysis is normally focused on those neurons showing enhancement of multisensory responses as compared to their unimodal responses. However, in our dataset, we observed significant proportion of neurons with sub-additive multisensory effects. In order to simplify the comparison of inverse effectiveness between the different populations, we plotted our data in a semi-log space and fitted separate regressions for each functional sub-population (Fig. 4G). This revealed significant correlations resembling inverse effectiveness. While supra-additive units showed weak, positive inverse effectiveness (r = -0.30, p = 7.7×10^−5^), sub-additive and additive units exhibited negative inverse effectiveness (r = 0.60, p = 1.2×10^−28^ and r = 0.24 p = 0.05, respectively), i.e. an increase of multisensory suppression with decreasing unisensory response strength.

These results indicate that multimodal units show the largest non-linearity in summation at low stimulus saliency levels, both for enhancement and suppression. This mirrors the findings from unit recruitment (Fig. 3G-I) and the behavioral detection task (Fig. 1G), where the largest changes are observed at low stimulus levels. Interestingly, although we observed similar effects of multisensory additivity and the recruitment of units during multisensory stimulation under isoflurane anesthesia, inverse effectiveness correlations were weak and only significant in the sub-additive population (Fig. S2G). While prior work has mostly focused on the enhancement of multisensory responses, our results highlight that a significant aspect of cortical multisensory integration is the interplay between multisensory enhancement and suppression of neural responses.

### Thermotactile responses in fS1 are prolonged and include additional action potentials

Non-linear changes of responses in multisensory integration are usually reflected by a quantitative change in the absolute number of APs during the entire stimulus period. However, a neuron’s response can also be changed qualitatively by altering its temporal response dynamics. In our population, we observed 3 major types of unimodal response dynamics: cells with an excitatory transient response (e.g. Fig. 3A, Fig. 4B top), cells with an excitatory sustained response (e.g. Fig. 2D top left, Fig. 4B middle) and cells whose response was suppressed (e.g. Fig. 2D bottom right).

In order to investigate the relationship between response dynamics during multisensory stimulation and additivity further, we applied hierarchical clustering to cool and touch responses separately in the sub-additive, supra-additive and additive populations (with 3 target clusters for each response type, i.e. a total of 3 × 3 = 9 combinations, see Fig. S5A and Methods) and graphically ordered them by the response dynamic during multisensory stimulation (top, sustained; middle, transient; bottom, suppressed, Fig. 5A, B). While the additive population did not show any significant differences in their temporal dynamics between unisensory and multisensory stimulation (Fig. S5B), both the supra- and sub-additive populations exhibited significant differences. In the sub-additive population, the differences were most obvious with ‘suppressed’ units (Fig. 5A, e.g. cluster 9), which highlights the role of inhibition in this sub- population. In contrast, the supra-additive population contained many units that were unmasked when both stimuli were presented together (Fig. 5B, e.g. clusters 2 and 3)., Strikingly, these clusters contained a large number of units with more sustained responses to multimodal than unimodal stimulation, indicating that response prolongation could be a central aspect of thermotactile integration.

**Figure 5:**
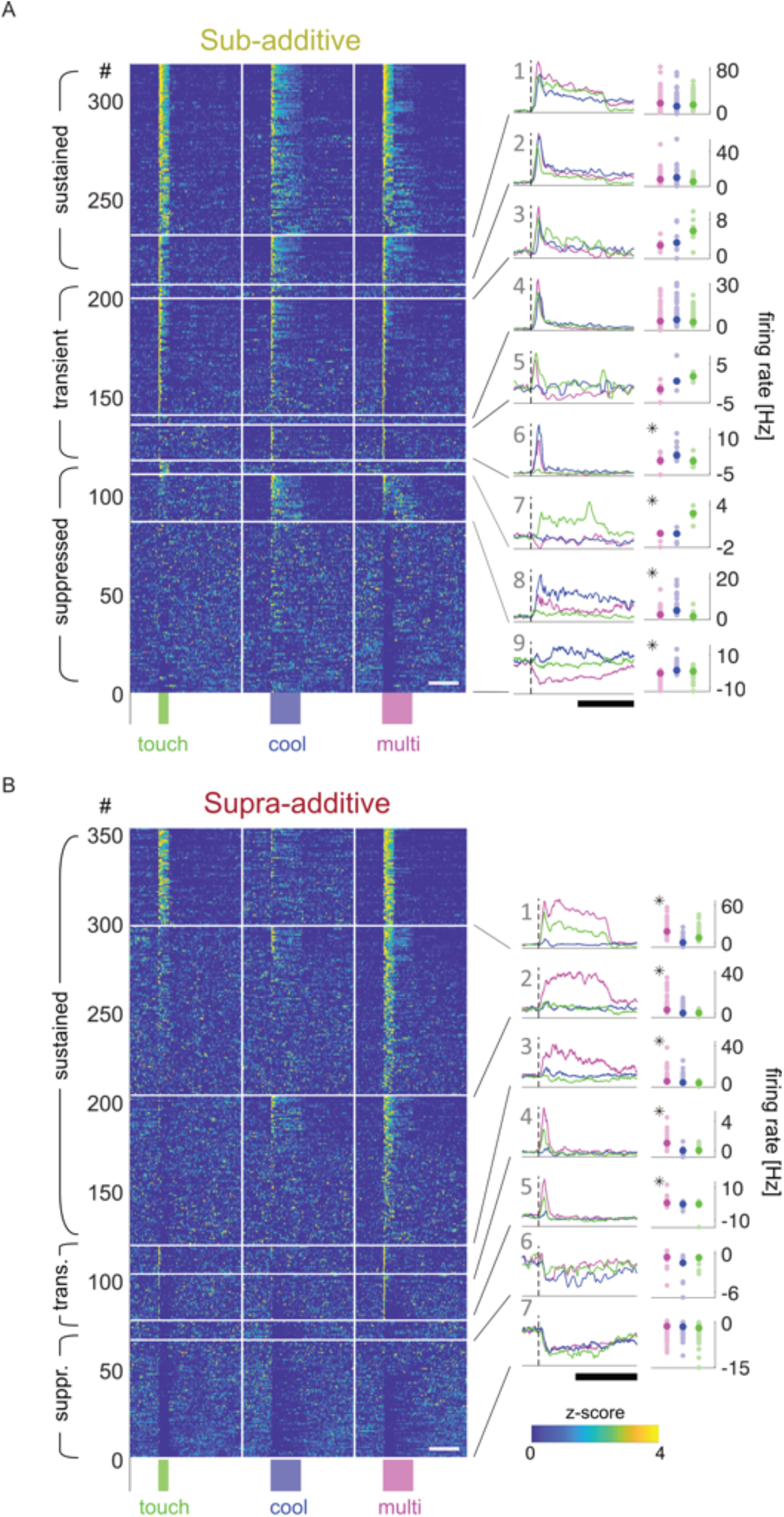
Temporal response dynamics of supra-additive units change during multimodal stimulation. A). Hierarchical 3-by-3 clustering of ‘best’ multimodal responses of the sub-additive sub-population. Each row shows the concatenated, z-scored responses to touch (left), cool (middle) and multimodal (right) stimulation for each unit. Clusters are ordered by temporal dynamics of the average multimodal response (left side panels) from sustained over transient to no response/suppressed. Within each cluster, units are ordered by peak response strength in the multimodal condition. Right side panels show the median responses of each cluster, asterisks denote significant differences between the multimodal and the best unimodal response (p<0.01, Wilcoxon test). X-axis scale bars are 3 seconds (white) and 500 ms (black). B). Same as in A) but for the supra-additive sub-population.

To examine this further, we first averaged and normalized the multimodal responses across the sub-additive, supra-additive and additive populations (Fig. 6A, left). Visual inspection shows a more prolonged multisensory response for the supra-additive neurons as compared to the sub-additive and additive populations across the entire population. To quantify this, we calculated a ‘duration index’ for each unit that compared the strength of the first and last 200ms of the response (Fig. 6B, left). We found that the distribution for supra-additive units was significantly shifted to the left compared to sub-additive units, confirming a longer response duration during multisensory stimulation, indicating this response feature is specific to the integration of thermotactile information (Fig. 6A, B, right panels).

**Figure 6:**
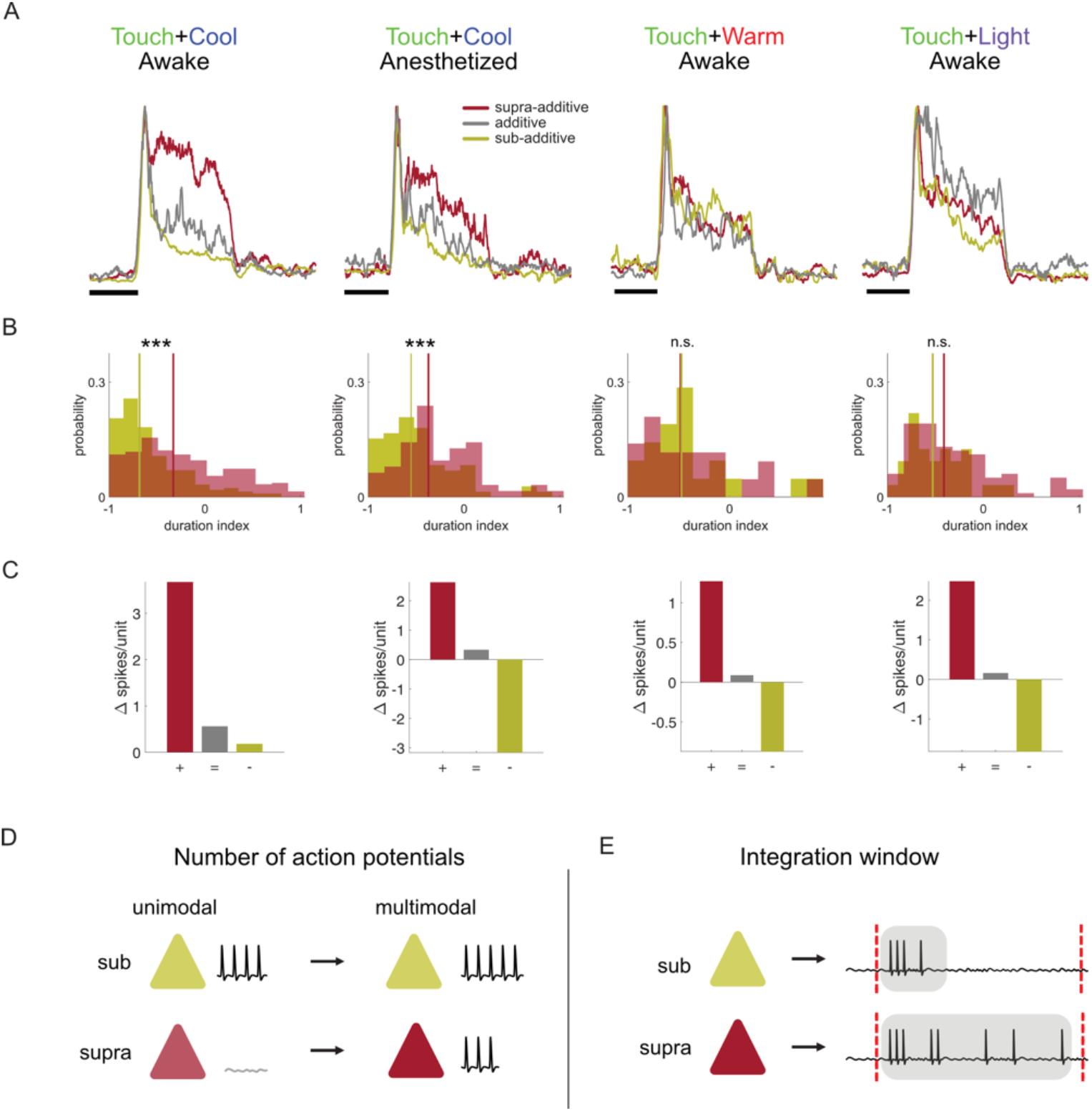
Supra-additive units strongly influence the population representation of thermo-tactile stimuli. A) Normalized, average response of all sub-populations. The supra-additive population exhibits more sustained response dynamics for cool/touch integration, an effect that is lost for warm and light control conditions. B) The duration index DI = ((L-E)/(L+E)) compares the 1st (E) and last (L) 200 ms of the response to generate a measure for ‘sustained’ and ‘transient’ responses. For touch/cool integration, the duration index for multimodal responses from supra-additive units is shifted to the right compared to sub-additive units, indicating ‘prolongation’ of the response. Lines denote medians of the respective distributions (Wilcoxon test from left to right: p = 4.7×10-15, p = 8.0×10-4, p = 0.13, p = 0.88). C) Average number of spikes per unit generated or lost in the multimodal condition by calculating the absolute difference in spikes/unit between the multimodal and best unimodal response across all clusters of each sub-population. Only in the awake cool/touch experiment there is a bias towards more spikes in the network while in the anesthetized and control experiments the number of added/lost spikes is more balanced. D) The number of action potentials generated in the multimodal vs. unimodal condition is only slightly increased (or even decreased) in the sub-additive population (yellow, ‘ceiling effect’) whereas there is a large, relative increase in the supra-additive population (red). E) The effective integration time window (shaded area) is longer in the supra-additive population due to sustained response dynamics. Red dotted lines denote stimulus window.

A change from more transient to more sustained response dynamics may result from the same number of APs being more distributed in time, or an absolute increase in the total number of APs. To address this question, we calculated the number of additional APs per unit in the multimodal versus the best unimodal condition for each sub-population (Fig. 6C). This analysis showed that the supra-additive population added APs/unit as compared to the sub and additive populations (Fig. 6C, left panel) and acted to boost the number of APs across the entire population during multisensory stimulation. This was also observed in the supra-additive population under anesthesia, but, in contrast to thermotactile stimulation in awake mice, there is a reduction in the number of APs in the sub-additive population during touch/cool stimulation in anesthetized mice and during touch/warm and touch/light integration stimulation in awake mice which counteracts the increase in numbers of APs in the additive population (Fig. 6C, right panels). Together these data suggests that a prolongation of the integration window in fS1 is a key property of thermotactile integration (Fig. 6D,E).

## Discussion

Somatosensation involves a continuous integration of temperature and tactile information. Here we examined the impact of integration of cool and tactile information on perception and neural representation in mouse forelimb primary somatosensory cortex (fS1). We show that perception is enhanced by the addition of a second modality, and pinpoint key features of cortical neural responses that are relevant for thermotactile perception; including, (i) the recruitment of unimodally silent or inhibited neurons, (ii) non-linearity of thermotactile responses, (iii) a prolongation of response duration.

### Thermotactile perception

To examine the impact of thermotactile integration on detection sensitivity (a change in perceptual threshold) and reliability (a change in the slope of the psychometric function), we designed a thermotactile Go/NoGo detection task based on an audio-visual integration task for head-fixed mice ^24,37^. Prior work suggests that multisensory modulation is enhanced if two stimuli are presented at the same environmental location (spatial congruency), the same time (temporal congruency) and at threshold stimulus levels (inverse effectiveness). To achieve spatial congruency, we delivered both stimuli to the right forepaw. However, the touch stimulus was delivered by vibrating the entire forepaw via the back of the paw, whereas the thermal stimulus was delivered to the glabrous skin on the bottom of the paw. Despite this limitation, both cool and touch stimuli evoked strong and reliable responses in fS1, demonstrating overlapping receptive fields in many neurons. To maintain temporal congruency and mimic the simultaneous timing of cool and touch stimuli during active object touch, we delivered both stimuli at the same onset times during thermotactile stimulation. Our task contained multiple amplitudes of both stimuli as well as an adaptive staircase procedure for session-by-session calibration of sensory threshold of both modalities to allow pooling of data across sessions, fitting of psychometric functions and an assessment of inverse effectiveness.

Our data show that mice show more accurate reporting at higher stimulus amplitudes and have low unimodal cool and touch detection thresholds (cool ∼0.5 °C, touch ∼2 mN). Moreover, we observed a longer detection latency to cool (245 ms) than to touch (219 ms) stimuli. It is challenging to compare perceptual features across modalities, but a likely explanation could be the established differences in neural conduction time for thermal and tactile information to travel from the skin sensory afferent neurons to the cortex (Fig. 3D). Cool sensitive primary sensory afferent neurons are thought to contain a population of slowly conducting c-fibers, as well as a smaller population of faster conducting, thinly myelinated primary thermal afferent neurons ^38–40^, whereas the tactile pathway contains a major population of fast conducting, myelinated afferent fibers and is optimized for fast transfer of information ^41^.

The central result from multisensory perceptual testing was that the detection of combined thermotactile stimuli was enhanced compared to unimodal stimuli. This was shown by an increase in the detection rates (Fig. 1C) and a reduction in the detection time compared to unimodal stimuli (Fig. 1E). Notably, the difference between maximum unimodal and bimodal detection rates (Fig. 1G) was stronger at threshold stimulus levels than at higher or lower amplitudes. Further analysis of the psychometric functions showed a significant reduction in perceptual thresholds during multisensory stimulation (Fig. 1D), but, consistent with prior studies of audio-visual detection ^42,43^ but see ^37^, there was no overall change in the slope of the psychometric function during thermotactile stimulation. Therefore, the enhancement of task performance was likely the result of the reduction in perceptual threshold (Fig. 1D) rather than an increase in detection reliability.

Could these results be explained by the mouse being poorly motivated or using limited information in unimodal compared to multimodal testing? This seems unlikely because we observed that the shape of the response time distribution was similar across all conditions (Fig. 1E), suggesting similar motivation and similar use of available sensory information during uni- and multi-modal testing ^5^. Instead, we hypothesized that behavioral enhancement results from neural response features of cortical neurons during thermotactile integration. To test this, we went on to perform extracellular recordings from fS1 neurons in resting, awake mice.

### Cortical representation of thermotactile integration

Prior work has shown that mouse fS1 neurons respond to cool and touch stimulation of the forepaw ^33,34^, but their thermotactile integration properties had not been investigated. In this study, we aimed to examine cortical mechanisms of thermotactile integration and, to avoid non-sensory input to fS1, we did not combine electrophysiological recordings with behavioral testing. In the future, two alternative forced choice paradigms with delay periods could help address these issues.

We confirmed that fS1 neurons show robust responses to unisensory cool and touch stimuli (Fig. 2D) and that many neurons encode both modalities. Similar to the enhanced behavioral reporting of multisensory stimuli, the overall cortical response strength was higher for multisensory than unisensory stimuli (Fig. 3B). In line with the unimodal behavioral response latencies, cortical responses to touch stimuli had shorter latencies than to cool stimuli, but in contrast to the reduction in thermotactile perceptual reporting times, the mean neural latency to thermotactile stimuli was in between the touch and cool response latencies, possibly reflecting the fact that multisensory unit’s responses can be dominated by either modality.

An interesting finding was that many neurons with no change or even a reduction in firing rates during unimodal stimulation showed significant responses during multisensory, thermotactile stimulation (Fig. 5B clusters 2,3). This boosting of responses could be a key feature of multimodal stimulus representation in fS1 (Fig. 3I). One possibility is that the recruitment of previously silent cells could result from alterations in the level of local synaptic inhibition during multisensory stimulation ^22,44^. Future experiments could address this hypothesis with whole-cell, membrane potential recordings from cortical neurons combined with activity manipulations of cortical GABA-ergic inhibitory interneurons.

A central question regarding cortical multisensory integration is whether stimuli are combined in a linear or non-linear fashion. We tested this by comparing the linear addition of unimodal responses with the combined, multisensory response. Across the entire population of neurons responding to at least one modality, we found evidence for different forms of integration including non-linear supra- and sub- additive responses as well as linear additive responses (Fig. 4, Fig. S5B). Plotting a standard measure of multisensory responsiveness, the ‘Multisensory Index’, against the sum of the unimodal responses showed opposing forms of inverse effectiveness for the functional subtypes. As the sum of the unimodal responses decreased, the Multisensory Index of supra-additive neurons increased which would result in a boost of low-saliency inputs in this populations. Conversely, the Multisensory Index of the sub-additive population decreased, which could be interpreted as a suppression of noisy inputs in an already strongly responding population. Although inverse effectiveness is a well-documented form of non-linear integration and has been proposed as a key mechanism of multisensory integration in several systems.^25,37,45–47^ it is not straightforward to assess due to caveats including the ‘ceiling/floor effects’ and ‘regression towards the mean’ ^48,49^. Our stimulation paradigm used an a priori approach which samples responses to a range of defined stimulation intensities, as opposed to sorting neuronal responses by strength. This may mitigate problems associated with ‘regression towards the mean’ ^49^, but issues with noisy estimations of small responses (floor effect) and finite maximum firing rates of neurons (ceiling effect) remain. This makes it difficult to unambiguously identify inverse effectiveness as a key mechanism of multisensory enhancement in our recordings.

At the same time, we observed a clear difference between the response dynamics of supra-additive, sub-additive and additive neurons to thermotactile stimulation. The supra-additive neurons showed a sustained responses during thermotactile stimulation (e.g. Fig. 3A, 4B), which was not the case in the sub-additive or additive populations. A prolongation of responses especially at low saliency levels could favor a more robust representation over time for hard-to-detect stimuli by increasing the effective integration time window. We also observed that the prolonged responses lead to additional spikes in the supra-additive population (Fig. 6C). Intriguingly, the subset of neurons with the clearest response prolongation were only weakly responsive or even silent during unimodal stimulation (Fig. 5B, clusters 2 and 3), suggesting that they are being recruited in the multimodal condition (Fig. 3I, Fig. 6D, E). Together with the observed delay in response latency at low saliency levels during unisensory perception (Fig. S1B), the prolongation of the cortical response could be a substrate for the performance increase during multisensory perception.

Cortical sensory processing is also known to be modulated by arousal and differences in stimulus saliency or behavioral engagement. Even in resting, awake mice, arousal systems could be activated to different extents and contributed to the multisensory effects we observed in fS1. To assess the role of arousal, we performed recordings under anesthesia, where arousal levels are more stable than in awake mice. Interestingly, inverse effectiveness was only present in the sub-additive population under anesthesia, whereas response prolongation and unit recruitment effects were comparable to awake mice. As a control, we went on to compare cool/touch responses with the responses to multisensory stimuli that are not well represented in fS1 (warm/touch and light/touch). These results showed that the prolongation of responses and the population level increase in spiking was present only during cool/touch stimulation, suggesting that they are the result of integration between two modalities of sensory input rather than generalized arousal.

### Conclusions and outlook

In conclusion, our behavioral results show that the integration of cool and touch inputs acts to enhance somatosensory perception. They suggest that future studies should consider the temperature of surfaces as a fundamental component of tactile sensation and attempt to dissociate thermal from tactile information ^50^.

Our electrophysiological recordings establish mouse fS1 as a model cortical region for investigations into the neural mechanisms of thermotactile integration. The recruitment of additional cortical neurons, non-linear changes in responses and longer response duration are likely key changes in fS1 that underlie the enhancement in perception during thermotactile integration (Fig. 3I, Fig. 6D,E). Temporally precise neuronal activity manipulations during a thermotactile integration tasks will be required to test this hypothesis.

## Acknowledgements

This work was supported by the European Research Council (ERC-2015-CoG-682422, J.F.A.P.), the Deutsche Forschungsgemeinschaft (DFG, FOR 2143, J.F.A.P., SFB 1315, J.F.A.P.), the German Academic Exchange Service (DAAD, I.E.-R.), Experimental Psychology Society (EPS, I.E.-R.) and the Helmholtz Society (J.F.A.P.).

