## Supplementary Information for "Cortical cellular encoding of thermotactile integration"

**Supplemental information contains:**

Methods

Supplemental Figures S1 – S5

### METHODS

#### Animals

All experiments were conducted according to European law and the state of Berlin animal welfare body (LaGeSo). Male C57BL/6J mice between P52 - P198 (median: P68) were used and maintained on a 12:12 hr light-dark cycle. Experiments were performed during the light phase. Mice were gradually habituated to head and paw fixation.

#### Surgery

Mice were deeply anesthetized using 3-4 % isoflurane in 100 % $O_2$ (maintained at 1.5-2 % isoflurane) and injected with metamizol (200 mg / kg) and 0.3-0.5ml warm sterile saline solution to avoid post-operative pain and ensure hydration. Mice were fixed in a stereotactic frame (SR-9AH, Narishige, Tokyo, Japan) using ear bars, eye gel (Vidisic, Bausch+Lomb, Laval, Canada) was applied and body temperature was maintained using a heating pad and rectal probe (50300, Stoelting, Wood Dale IL, USA). Custom-made head holders were attached to the cleaned skull and a well was build using dental cement and covered with KwikCast (World Precision Instruments, Sarasota FL, USA). After surgery, mice were kept on the warm heating pad until they awoke from anesthesia. Metamizol was dissolved in the drinking water for minimum 48 hr to avoid post-operative pain.

#### Intrinsic imaging

On the day of recording, mice were deeply anesthetized and then maintained at light anesthesia levels (1-1.5 % isoflurane) for IOI. The forepaw was stimulated with a 100 Hz sinusoidal vibration for 5 s using a Piezo element (PL127.11, Physik Instrumente, Karlsruhe, Germany), image frames were captured with a CMOS camera (QICAM Fast 1394, Teledyne, Surrey, Canada) and analyzed online using custom software written in Igor (WaveMetrics, Lake Oswego OR, USA) or MATLAB (Mathworks, Natick CA, USA). Once a robust response was detected, a craniotomy was performed over the area and 1 or 2 incisions were made in the dura to facilitate probe insertion. For awake recordings, the craniotomy was then covered with a drop of Ringer's and the well was sealed with KwikCast. The animal was allowed to wake up and recover from anesthesia for at least 1h before starting the extracellular recording procedure.

**Sensory stimulation**

Stimulus presentation was controlled via custom-written MATLAB-scripts and a DAQ-board (NI-6232, National Instruments, Austin TX, USA) and synchronized with the recording setup (see below) via TTL pulses. Additionally for the behavior experiments (see below), the trial structure and real-time feedback was controlled with a state machine (bpod, Sanworks, Rochester NY, USA).

Touch stimuli were delivered via a force-feedback lever system (300C, Aurora Scientific, Aurora ON, Canada) as a sinusoidal vibration to the top of the paw for 1 s at 60 Hz at different intensities. The force at the tip of the lever was recorded to detect movement of the animal during the recording and to implement an online feedback correction to keep the stimulus intensity consistent across trials in case the animal slightly moved the paw against the tape fixation.

Temperature stimuli were delivered via 5 high-performance 3.2 x 2.4 mm Peltier elements (TCSII, QST Labs, Strasbourg, France) covering the whole paw of the animal. The stimulus consisted of a ramp-hold-return of (1 s duration for behavior and 3 s duration for electrophysiological recordings) at a ramp speed of 300 °C / s covering a range of +/-8 °C from baseline (32 °C) at a resolution of 0.1 °C.

Visual stimuli were delivered via a white LED positioned ~8 cm from the contralateral eye mimicking the ramp-hold-return temperature stimulus at intensities of 12.5, 25, 50 and 100 cd/m$^2$ (calibrated using ColorCAL MkII, Cambridge Research Systems, UK). In a subset of experiments, a second Neuropixels probe was inserted into primary visual cortex V1 to ensure that the visual stimulation was eliciting primary sensory responses (data not shown).

**Behavioral task**

The multisensory Go / NoGo detection task and analysis was modelled after Meijer et al., 2018. Animals (n = 8) were trained in several steps to detect both cool and touch stimuli until they surpassed a d’ criterion of 1.5 for each step: 1) stimulus pairing 2) blocks for C or T only 3) C and T interleaved. We did not reward correct rejections and false alarms were not punished. In the testing sessions, psychometric curves for both modalities were generated separately by using an adaptive staircase method (PsychStairCase of the Psychophysics Toolbox for MATLAB, Brainard ‘97) in a Go/NoGo task (Fig. S1A). Multimodal trials were realized by combining the last used unimodal stimulus intensities from each staircase. All trials were pseudo-randomly interleaved (not more than 4x the same modality in a row). Animals performed on average 1959 trials (range: 1138-2944) over 5-11 sessions. Animals performing Go/NoGo tasks, can devise alternative response strategies to obtain rewards even at low performance levels or become over-motivated and generate false positives (Berditchevskaia, 2016). We therefore excluded periods where the FA-rate was above 20 % and used randomized periods (ITI 3-7 s + 4-5 s timeout) after each trial to discourage the animals from developing stereotypical licking patterns. Periods where the false alarm (FA) rate stayed on average over 20% were identified using a 100 trial wide sliding window and excluded from analysis.

#### Behavioral data analysis

Behavior data was analyzed using the Palamedes Toolbox (Prins and Kingdom, 2018). In short, maximum likelihood fitting with a Weibull function with 3 free (FA-rate, threshold, slope) and 1 fixed parameter (lapse rate) was used to estimate the psychometric functions for each condition. Lapse and FA rates were determined from blank and maximum intensity trials, respectively. Significant changes in threshold and slope between conditions were determined by comparing the transformed likelihood ratios of the full model to a model where either slope or threshold were fixed. The multimodal psychometric data ($cool_{touch}$ / $touch_{cool}$) was created by using all multimodal trials where the 'other' modality's intensity was subthreshold with respect to its unimodal psychometric curve. Subjective intensity was calculated by binning trials according to their stimulus intensity into an equal 5 x 5 grid and then calculating a weighted average of the detection rate across mice according to the number of trials in each bin per mouse. Multisensory enhancement was calculated as the % difference between the multimodal and best (i.e. highest) unimodal response rate for a given bin.

#### Extracellular recordings

All recordings were performed with Neuropixels1.0 probes (imec, Leuven, Belgium) and the spikeGLX acquisition package (Bill Karsh, Janelia Research Campus, Ashburn VA, USA) via a PXI interface (NI-PXIe-1071, National Instruments, Austin TX, USA). Probes were coated with DiI (Invitrogen, Waltham MA, USA) and lowered into the tissue automatically via micromanipulators (LN25, Luigs&Neumann, Ratingen, Germany) to a recording depth of 1500-1800 µm at a speed of ~2 µm / s. Raw data was acquired at ~30 kHz (actual rate calibrated for each probe separately) from up to 384 electrodes.

#### Histology

After the recording, mice were decapitated under deep anesthesia and the brain was immersed in paraformaldehyde (PFA) for minimum of 24 hr. 100 µm thick slices were cut on a vibratome (VT12000S, Leica, Wetzlar, Germany) and imaged on a fluorescence microscope

(AxioImager.M2, Karl Zeiss Microscopy GmbH, Göttingen, Germany) to visualize the DiI probe track. Slice images were subsequently transformed and aligned to the Allen Institute mouse brain atlas using the SHARP-track package (Shamash et al., 2018) to confirm the location of the recording in fS1.

**Extracellular spike sorting**

Spike sorting was performed offline using Kilosort 2.0 (Pachitariu et al., 2016). Manual curation of the sorted units was performed using the Phy2 package (https://github.com/cortex-lab/phy). Splitting and merging of clusters was kept to a minimum and manual curation was mostly used to tag putative single units and exclude drifting units and artifacts for subsequent analysis. After manual curation, all remaining units were evaluated by 3 main quality metrics: minimum number of spikes > 1000, refractory period (1.5 ms) violations < 0.5%, number of missing spikes < 10% (evaluated as the overlap of 2 gaussian fits to the waveform and noise amplitude distributions).

**Neuronal analysis**

Offline analysis of stimulus responses was performed for each putative single unit using custom MATLAB scripts and existing code (https://github.com/cortex-lab/spikes). Responses were defined as the average, baseline-corrected firing rate (or z-score) in response windows of varying size (default: 1 s) after stimulus onset. The stimulus space consisted of 4 x 5 = 20 amplitude combinations for touch and temperature (0, 5, 10, 20 mN and 0, ±1, ±2, ±4, ±8 °C) with 25 repetitions per stimulus. Units were considered 'responsive' to a particular stimulus if the number of APs measured in the response window deviated significantly from the spiking probability of a Poisson process with that unit's mean baseline firing rate by using a 'binless' method (i.e. using 'bins' of 1ms, $\alpha < 0.01$, Antoine et al, 2019). The first 'bin' to cross the significance threshold of $\alpha < 0.01$ was used as the latency of this unit's response. Units were tagged as 'overall responsive' if at least 2+x out of 20 stimuli elicited significant responses with x being the rounded false positive detection rate of the binless method determined from blank trials multiplied by 20 (average over 22 awake recordings: x = 5.12 ± 0.47 %; average over 6 anesthetized recordings: x = 10.22 ± 2.91 %).

PSTHs were constructed using an optimal binsize algorithm (Shimazaki and Shinomoto, 2007). Instantaneous firing rate was calculated using a gaussian KDE. Hierarchical clustering was performed on the z-scored KDE over the whole trial of a particular stimulus combination (maximum unimodal stimulus intensities (20 mN / -8°C) for 2 x 2 clustering; 'best' stimulus combination from additivity analysis for 3 x 3 clustering, see below) for each unit using the 'linkage' and 'cluster' functions in MATLAB.

The additivity of each unit was determined as follows: First, for each possible stimulus combination, the distribution of the arithmetic sum of the unimodal response values was bootstrapped (10000 iterations, 12 stimulus combinations) and then z-scored. If the z-score of the actual multimodal response surpassed ± 1.96, this response was considered supra- or sub-additive, respectively and additive otherwise). The stimulus combination which had the maximum absolute z-score was tagged as the 'best' combination. The Multisensory Index was calculated as the difference between the multimodal and the summed unimodal responses normalized by the sum of both (MSI = M - (T + C) / M + (T + C)).

**Statistics**

Statistical tests were performed using MATLAB built-in functions. Between two conditions (median/mean): Wilcoxon ranksum / paired t-test; Across several conditions: ANOVA (Kruskal-Wallis for multiple comparisons with Dunn-Sidak correction). If not otherwise noted, values are reported as mean ± SEM (standard error of the mean). P-values denoted by stars: ***: $p < 0.001$, **: $p < 0.01$, *: $p < 0.5$, n.s.: not significant.

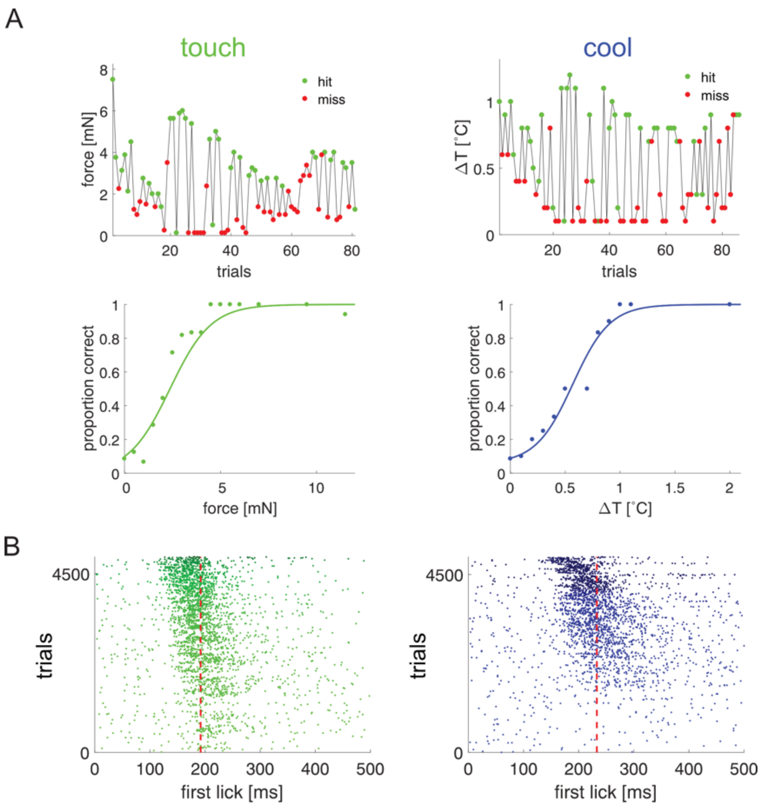


**Figure S1: Behavioral staircase procedure and licking latency**

A) Top, example stimulus detection during staircase testing for 1 session of an example animal for both touch and cool stimuli, each dot represents one stimulus trial. Bottom shows corresponding psychometric curve fits with each dot showing mean detection value for binned stimulus amplitude.

B) Lick latency correlates with stimulus intensity. Left shows all trials across all mice for touch (left) and cool (right) unimodal stimuli, ordered by stimulus intensity. Red dotted line denotes median first lick latency.

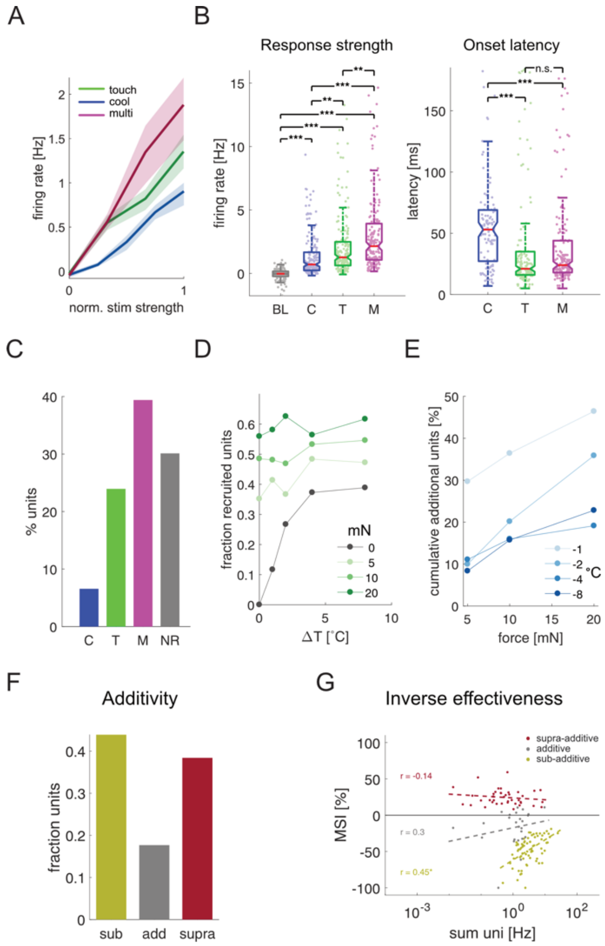


**Figure S2: Recruitment and additivity in multisensory integration are preserved under isoflurane anesthesia**

A) Average firing rate of all responsive units to increasing stimulus intensity for touch, cool and multimodal stimulation.

B) Median response strength (left) and onset latency (right) for all responsive units for all stimulus conditions (BL = baseline, C = cool, T = touch, M = multimodal). Significant differences were determined by multiple comparison testing (Kruskal-Wallis test).

C) Fractions of responsive (cool, touch, multimodal) and unresponsive single units (n = 259 total) at maximum intensity stimulation.

D) The fraction of recruited units as a function of stimulus intensity.

E) Same as in D) but for cumulative recruited units.

F) Fraction of units in respective sub-populations after determining additivity.

G) MSI plotted against the arithmetic sum of the corresponding unimodal stimulus response. Inverse effectiveness is estimated by linear fits to the different sub-populations of neurons (r =

0.3, p = 0.14; r = -0.14, p = 0.37; r = 0.45, p = 0.0001 for the additive and supra-/sub-additive populations, respectively).

A

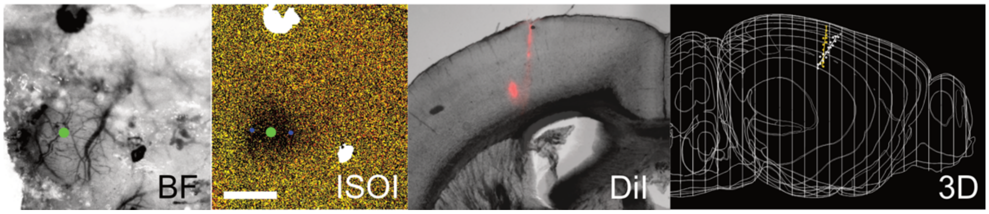


**Figure S3: Identification of forepaw S1 and tracking of recording electrode location**

A) Panels from left to right: (i) Craniotomy under brightfield (BF) illumination and (ii) corresponding intrinsic signal optical imaging (ISOI). Darker area denotes response to tactile stimulation of the forepaw. Scale bar is 1 mm. Green dot denotes the same spot in both panels. (iii) DiI-staining in a horizontal brain slice (100 μm thickness) of two Neuropixel probe tracts and (iv) 3-D reconstruction of both probe trajectories in a normalized mouse brain model (SHARP-track / Allen mouse brain atlas).

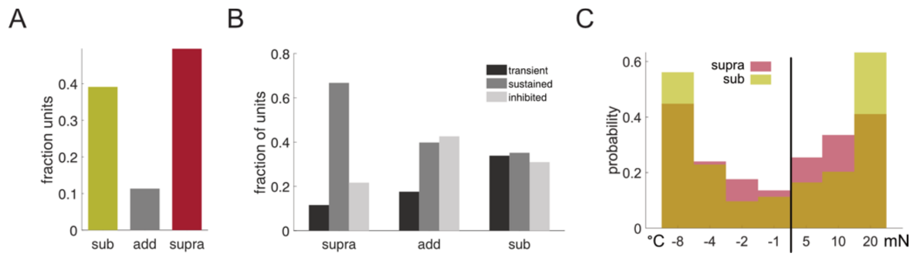


**Figure S4: Comparison of additivity sub-populations**

A) Fraction of excited units in respective sub-populations after additivity analysis.

B) Fraction of units that showed transient, sustained or suppressed response dynamics in each sub-population.

C) Distributions of stimulus combinations that produce 'best' responses in the multimodal condition for the sub-additive (yellow) and supra-additive (red) population.

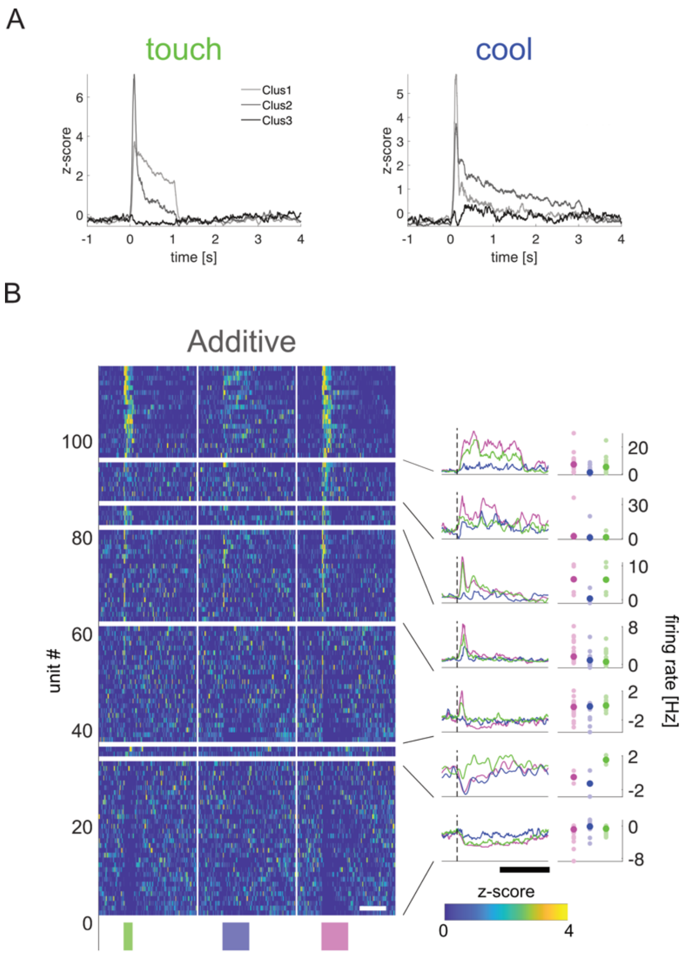


**Figure S5: Cluster temporal dynamics and clustering result of the additive sub-population**

A) Average z-scored responses of sub-clusters after hierarchical 3-by-3 clustering for touch (left) and cool (right).

B) Hierarchical 3-by-3 clustering of ‘best’ multimodal responses of the additive sub-population. Each row shows the concatenated, z-scored responses to touch (left), cool (middle) and multimodal (right) stimulation for each unit. Clusters are ordered by temporal dynamics of the average multimodal response (left side panels) from sustained over transient to no response/inhibited. Within each cluster, units are ordered by peak response strength in the multimodal condition. Right side panels show the median responses of each cluster. There were no significant differences between conditions. X-axis scale bars are 3 seconds (white) and 500 ms (black)
